# Intravital Two-Photon Microscopy of the Native Thymus

**DOI:** 10.1101/2023.04.10.536267

**Authors:** N. Seyedhassantehrani, C. S. Burns, R. Verrinder, V. Okafor, N. Abbasizadeh, J. A. Spencer

## Abstract

The thymus, a key organ involved in the adaptive immune system, is damaged by a variety of insults including cytotoxic preconditioning. This damage can lead to atrophy and potentially to changes in the hemodynamics of the thymic blood vascular system. Although the thymus has an innate ability to regenerate, the production of T cells relies on the trafficking of lymphoid progenitors from the bone marrow through the altered thymic blood vascular system. Our understanding of thymic blood vascular hemodynamics is limited due to technical challenges associated with accessing the native thymus in live mice. To overcome this challenge, we developed an intravital two-photon imaging method to visualize the native thymus in vivo and investigated functional changes to the vascular system following sublethal irradiation. We were able to quantify blood flow velocity and shear rate in cortical blood vessels and identified a subtle but significant increase in vessel diameter and barrier function ~24 hrs post-sublethal irradiation. Ex vivo whole organ imaging of optically cleared thymus lobes confirmed a disruption of the thymus vascular structure, resulting in an increase in blood vessel diameter and vessel area, and concurrent thymic shrinkage. This novel two-photon intravital imaging method enables a new paradigm for directly investigating the thymic microenvironment in vivo.

## Introduction

The thymus is a primary lymphoid organ essential for T cell development and a cornerstone of the adaptive immune system. T cell development relies on the complex interaction between thymus stromal cells (e.g., thymic epithelial cells (TECs), blood vascular endothelial cells (EC), and fibroblasts) and thymocytes which themselves are dependent on the recruitment of de novo “seeding” early thymic progenitor cells (ETPs) from the bone marrow (BM) (Seggewiss & Einsele, 2010; Velardi et al., 2013). The thymus is extremely sensitive to a range of acute and/or chronic insults, such as stress (corticosteroids), infection, sex hormones, and many cytoreductive treatments including chemotherapy, radiotherapy, and antibody therapy (Chaudhry et al., 2016). For example, the damage caused by cytoreductive treatments in patients undergoing hematopoietic cell transplantation (HCT) inhibits the capacity of the thymus to produce functional T cells contributing to increased morbidity and mortality (D’Souza et al., 2020; Krenger & Holländer, 2010; Schwarz & Bhandoola, 2004; Velardi et al., 2013; Zlotoff et al., 2011). Endogenous thymus regeneration after total body irradiation depends on the engraftment of BM lymphoid progenitors as ETPs in the thymus. They mobilize into the bloodstream from the BM and traffic to the thymus through the blood vascular system and enter the thymus at the cortico-medullary junction (Buono et al., 2016; Prockop & Petrie, 2000; S. W. Rossi et al., 2006; Zlotoff & Bhandoola, 2011). Early on, ETPs interact with thymic ECs to begin the differentiation process into distinct T-cell subsets (Buono et al., 2016; Mori et al., 2007; Ren et al., 2022; F. M. V. Rossi et al., 2005; Shi et al., 2016; Zhang & Bhandoola, 2014; Zlotoff et al., 2010). The thymus vasculature and ECs play critical roles as the highways of entry and source of cell signaling, respectively, during the early stages of endogenous thymic regeneration (Buono et al., 2016; F. M. V. Rossi et al., 2005; Zlotoff & Bhandoola, 2011).

It was previously reported that thymic ECs are radio-resistant and that the absolute number of ECs does not significantly decrease after sublethal total body irradiation (SL-TBI) (Wertheimer et al., 2018). Nevertheless, visualization of the thymic blood vessel network using light-sheet fluorescence microscopy 4 days after SL-TBI revealed that the total volume of the vasculature decreases, along with the total number of vessel segments, average vessel length, and vessel branching (Wertheimer et al., 2018). Notwithstanding the critical role of ECs in thymic regeneration, it remains unclear if these changes to the vascular network occur earlier than 4 days and whether these changes alter thymic hemodynamics or other vascular functions.

To study functional changes to the vasculature, high-speed intravital microscopy with subcellular resolution is required, but direct visualization of the native thymus in live mice has been elusive and deemed impossible with two-photon microscopy (Aghaallaei & Bajoghli, 2018). The position of the thymus directly underneath the sternum and internal thoracic vein, and adjacent to the heart and lungs, introduces several logistical, mechanical, and health complications when attempting to image the thymus. Standard methods such as flow cytometry, immunohistochemistry, and imaging of excised tissue have been applied to study the thymus in preclinical mouse models but these are limited to ex vivo analysis. Ex vivo culture systems (Hong & Moore, 1996; Ramsdell et al., 2006) including thymic slices (Bhakta et al., 2005; Ehrlich et al., 2009; Fournier et al., 2020; Le Borgne et al., 2009; Markert et al., 1997; Zhou et al., 2020) and whole organ imaging (Campinoti et al., 2020; Dzhagalov et al., 2012) are very powerful systems for studying certain aspects of thymus biology such as thymocyte-stromal interactions (Ladi et al., 2008; Nakagawa et al., 2012) and T cell development (Le Borgne et al., 2009; Witt et al., 2005), but these techniques lack blood flow and may not fully recapitulate the in vivo situation. Due to a lack of viable intravital imaging methods for the native thymus, researchers have relied on thymus transplantation to optically accessible sites including the kidney capsule (Caetano et al., 2012; Gregorio et al., 2021; Iwai & Inaba, 2015; Morillon et al., 2015), anterior chamber of the eye (Oltra & Caicedo, 2018), and pinna of the mouse ear (Chen et al., 2013) for intravital microscopy, but transplantation may alter the native vascular connections and exposes the tissue to an aberrant environment (Caetano et al., 2012; Vollmann et al., 2021). Although it has been reported that the transplanted thymus mirrors the endogenous thymus in architecture and vascularity, direct measurement of hemodynamic parameters in the native thymus is yet to be reported (Vollmann et al., 2021). Teleost fish have been suggested as a viable alternative to mice models since the native thymus can be directly observed in vivo (Bajoghli et al., 2015; He et al., 2020; Hess & Boehm, 2012; Li et al., 2007). However, in mammals such as mice and humans, T cell development critically and nonredudantly depends on the IL-7 cytokine signaling pathway which is not true in teleost fish (Lawir et al., 2019). Therefore, tools to directly visualize the native mouse thymus are needed in order to study functional changes to the thymic vascular system after cytotoxic preconditioning in an immunologically similar model to humans.

Here, we developed a novel intravital imaging method using two-photon microscopy to visualize the native thymus in live mice without transplantation. This method relies on thoracotomy to access the thymus within the chest cavity, followed by the placement of a custom-designed adhesion stabilization holder to reduce vertical and lateral movement from the heart and lungs (Aguirre et al., 2014; Jones et al., 2018; Lee et al., 2008, 2012, 2014; Vinegoni et al., 2015). Using this method, we were able to directly investigate functional and anatomical changes to the thymic blood vascular network within 24 hrs after SL-TBI in live mice. Two-photon intravital imaging of the native thymus opens a new paradigm for studying thymus biology which was not previously possible.

## Results

### Experimental Setup for Intravital Two-Photon Imaging of the Native Thymus

To image the native (i.e. in situ) murine thymus, we adapted methods for intravital cardiac microscopy (Lee et al., 2012; Vinegoni et al., 2015) to visualize and stabilize the thymus for intravital two-photon microscopy. Mice (8-12 weeks old) were anesthetized, ventilated, and put on a heated stage (**Fig. 1A)**. Next, a thoracotomy (Eichhorn et al., 2018) through the second intercostal space was performed to expose the thymus for imaging, and retractors were used to hold open the chest wall (**Fig. 1A, B**). To stabilize the tissue during intravital imaging, we designed and 3D printed a custom thymus adhesion stabilization holder to reduce movement artifacts caused by cardiac and respiratory motion (**Fig. 1C**). The stabilization holder consists of a flat ring at one end which is attached to a support arm mounted to the microscope stage. The underside of the ring was covered with a thin layer of Vetbond tissue adhesive (3M Vetbond), which bonds to the underlying thymic tissue and provides a stable field of view (FOV) for intravital two-photon microscopy (**Fig. 1D**). Proper placement of the stabilization holder helps to minimize vertical and lateral movement of the thymus and prevent blurring and blinking artifacts during imaging (**Supp. Fig. 1A, B**). Using this imaging setup, it was possible to have a clear imaging field within the thymus at a maximum diameter of 2.5 mm. To confirm tissue and vasculature viability in the thymus of untreated and SL-TBI (4.5 Gy) mice after preparing the thymus for imaging, Evans blue dye was injected retro-orbitally and visible perfusion of the thymus was observed in vivo and ex vivo (**Fig. 1E and Supp. Fig. 1C**).

**Figure 1.**
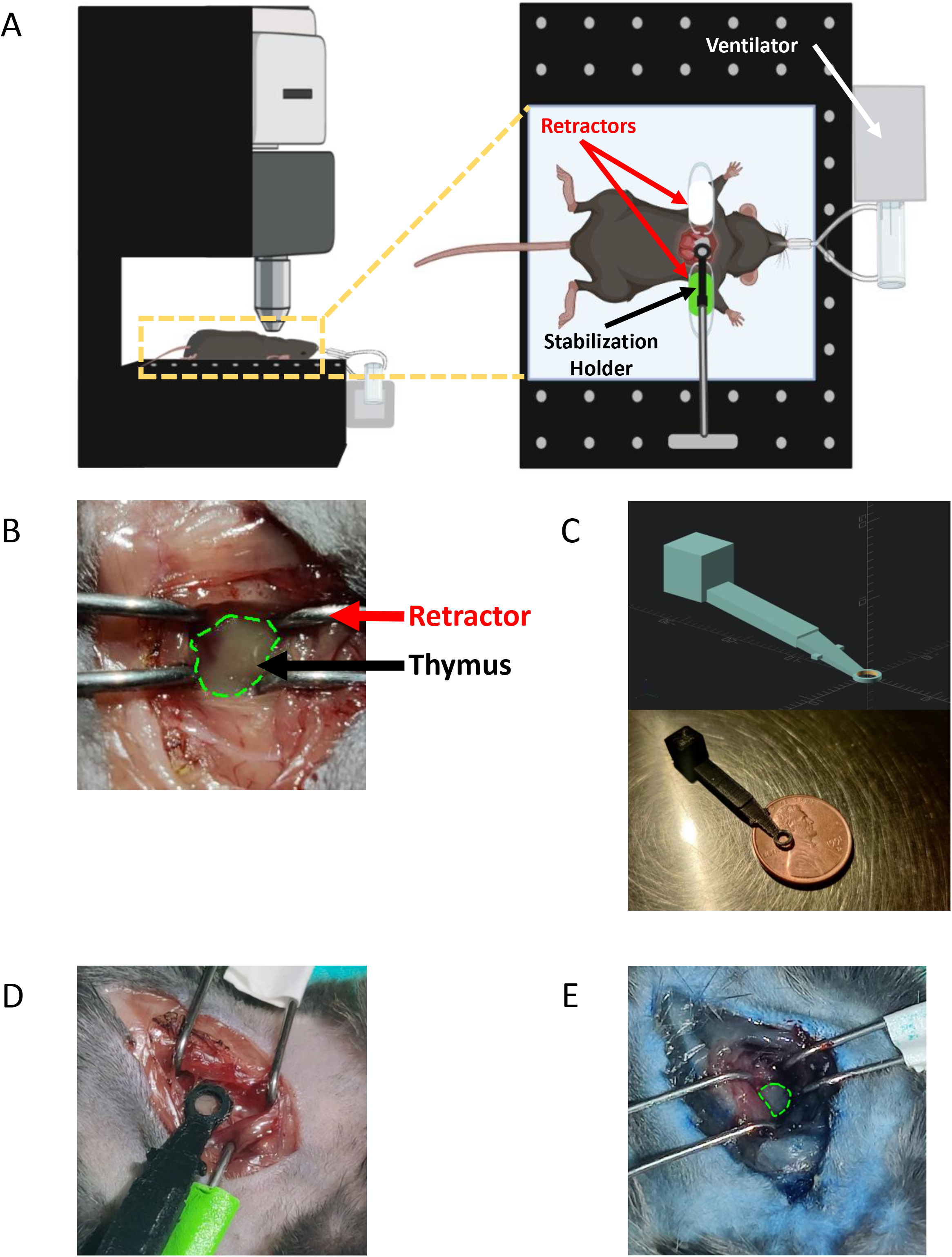
Experimental Setup and Thymus Holder Design. **(A)** Microscope diagram showing placement of chest retractors and adhesion holder around the thymus. **(B)** Thymus exposed through the 2^nd^ intercostal space. Green outline: Thymus. **(C)** 3D model of thymus adhesion stabilization holder and 3D-printed prototype. **(D)** Adhesion stabilization holder attached to the exposed thymus. **(E)** Perfused thymus after imaging indicates intact blood flow in the thymus at the time of Evans blue injection. Green outline: Thymus.

### In Vivo Imaging of the Thymus

We recorded two-photon images and videos in the native thymus of live mice using the methodology described above (**Fig. 2A-C and Supp. Movies 1-3;** n = 4 mice). Representative images and videos show the functional blood vessel network of the cortex and individual GFP-labeled cells within UBC-GFP mice (**Figs. 2A-C and Supp. Movies 1-3**). In these representative images and movies of the thymus, we can clearly see the thymus capsule, cells that compose the thymus microenvironment, and the blood vessel network. The stabilization adhesion holder minimized the motion generated by the heart and intubated lungs, allowing for optical sectioning and clear visualization of individual cells with subcelluar resolution within the thymus (**Fig. 2B and Supp. Fig. 1B**). In addition, we confirmed the presence of blood flow within individual blood vessels in the thymus via the movement of red blood cells (RBCs) visualized through negative contrast labeling (**Fig. 2C and Supp. Movies 2-3**). To confirm the structures observed during live imaging were from within the thymus, we dissected and imaged the thymus with two-photon microscopy ex vivo and detected very similar structures to the intravital images and movies (**Supp. Fig. 2A**).

**Figure 2.**
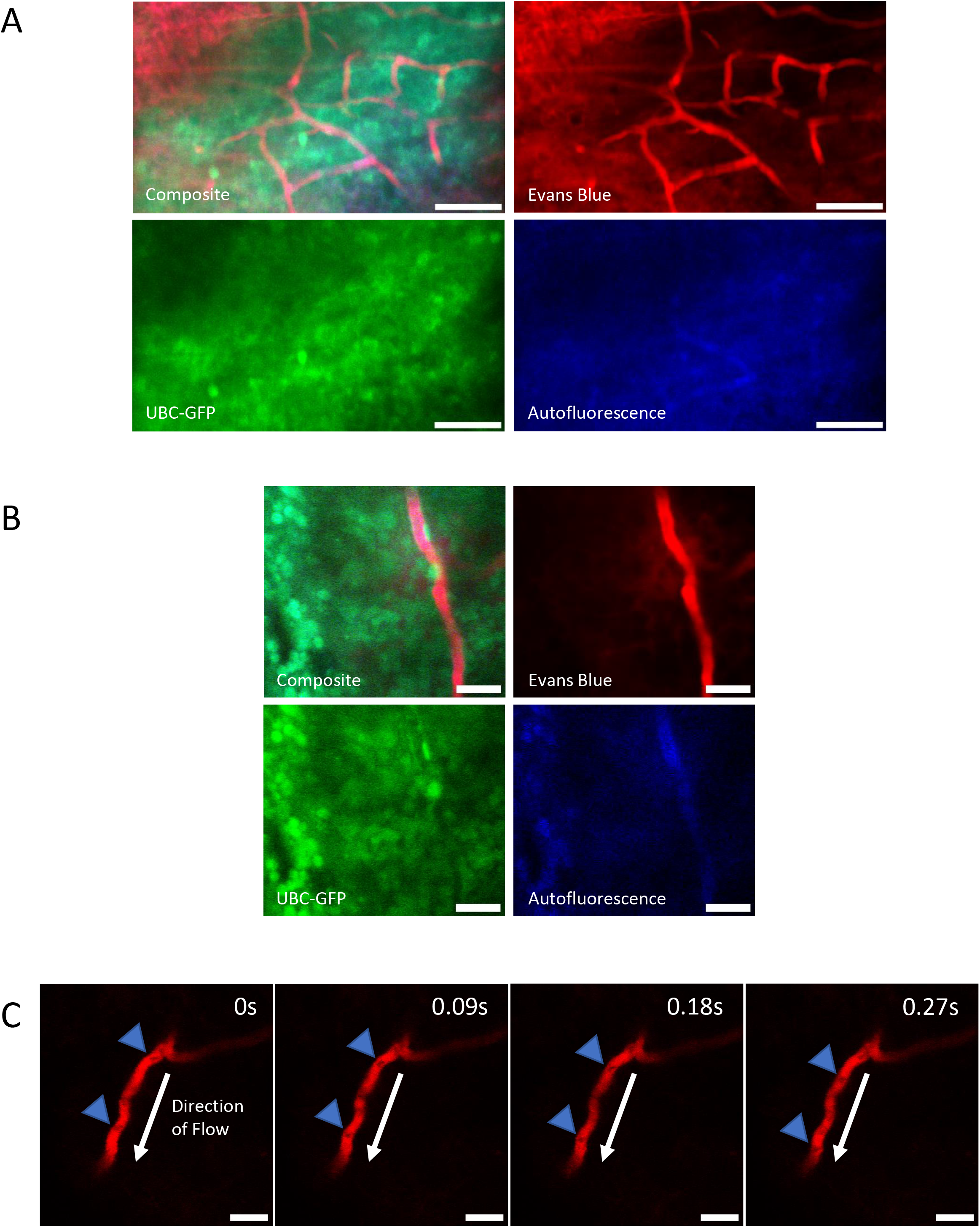
Intravital Imaging of Blood Flow in the Native Thymus. **(A**) Representative maximum intensity projection of the untreated thymus from a UBC-GFP mouse in vivo. Red: blood vessels (Evans Blue)/thymus capsule; Green: GFP; Blue: autofluorescence. (Scale bars ~ 50 μm). **(B)** Example average intensity projection of the untreated thymus from a UBC-GFP mouse demonstrating the ability to visualize individual GFP+ cells in the thymus in vivo. Red: blood vessels (Evans Blue); Green: GFP; Blue: autofluorescence. (Scale bars ~ 25 μm). **(C)** Example negative contrast labeled blood flow in the untreated thymus from a UBC-GFP mouse showing the movement of RBCs through blood vessels over time. Red: blood vessels (Evans Blue); Blue Arrow: negative contrast labeled RBC; White Arrow: direction of blood flow. (Scale bars ~ 25 μm).

### Effects of Irradiation on the Thymus Vasculature

Next, after imaging the native thymus, we hypothesized that changes to the blood vessel network structure from SL-TBI would alter hemodynamics, barrier function, and potentially lead to cessation of flow in some microvessels of the cortex. To investigate this hypothesis, we performed SL-TBI (4.5 Gy) on UBC-GFP mice (n = 3 mice), imaged them 24 hours afterward, and compared them to age-matched untreated UBC-GFP controls that were not given SL-TBI. Despite a significant reduction (difference = 27.7%, p = 0.0043) in the size of the thymus after SL-TBI (**Fig. 3A and Supp. Fig 2B**), which increased the difficulty of attaching the stabilization holder to the thymus, we were able to confirm the presence of blood flow within individual vessels in the irradiated thymus (**Fig. 3B and Supp. Movies 4-5**). We observed a nonsignificant difference (p = 0.1244) in thymic blood flow velocity between SL-TBI treated and untreated mice (mean velocity = 191.8 μm/s and 169.2 μm/s, respectively; **Fig. 3C**). In addition, a nonsignificant difference (p = 0.890) was observed in the shear rate between the SL-TBI treated and untreated mice (mean = 264.7 s^-1^ and 231.4 s^-1^, respectively; **Fig. 3D**). Nevertheless, we observed a small but statistically significant increase (p < 0.0001) in mean blood vessel diameter of SL-TBI treated vs. untreated mice (mean blood vessel diameter = 8.9 μm and 7.3 μm, respectively; **Fig. 3E**). Due to imaging depth limitations inherent to intravital two-photon microscopy, most vessels measured were likely capillaries within the thymus cortex. However, our observation of large diameter (>10 μm) blood vessels indicates that for some animals, it may be possible to image to the cortical-medullary boundary or even the medulla (**Fig. 3E**). We also observed the blood vessel barrier function was altered from SL-TBI resulting in significantly higher (p < 0.0001) vessel leakage in SL-TBI treated vs. untreated mice (mean leakage value = 0.4734 vs. 0.3278, respectively; **Fig. 3F**). Following intravital imaging, imaging of dissected thymi of SL-TBI mice revealed noticeable damage to the thymic microenvironment with widespread edema and an increase in autofluorescence **(Supp. Fig. 2C**). Overall, using our novel thymic imaging method, we demonstrated that the vascular barrier is compromised as early as 24 hrs after SL-TBI but that the blood flow is mostly unaltered at this time.

**Figure 3.**
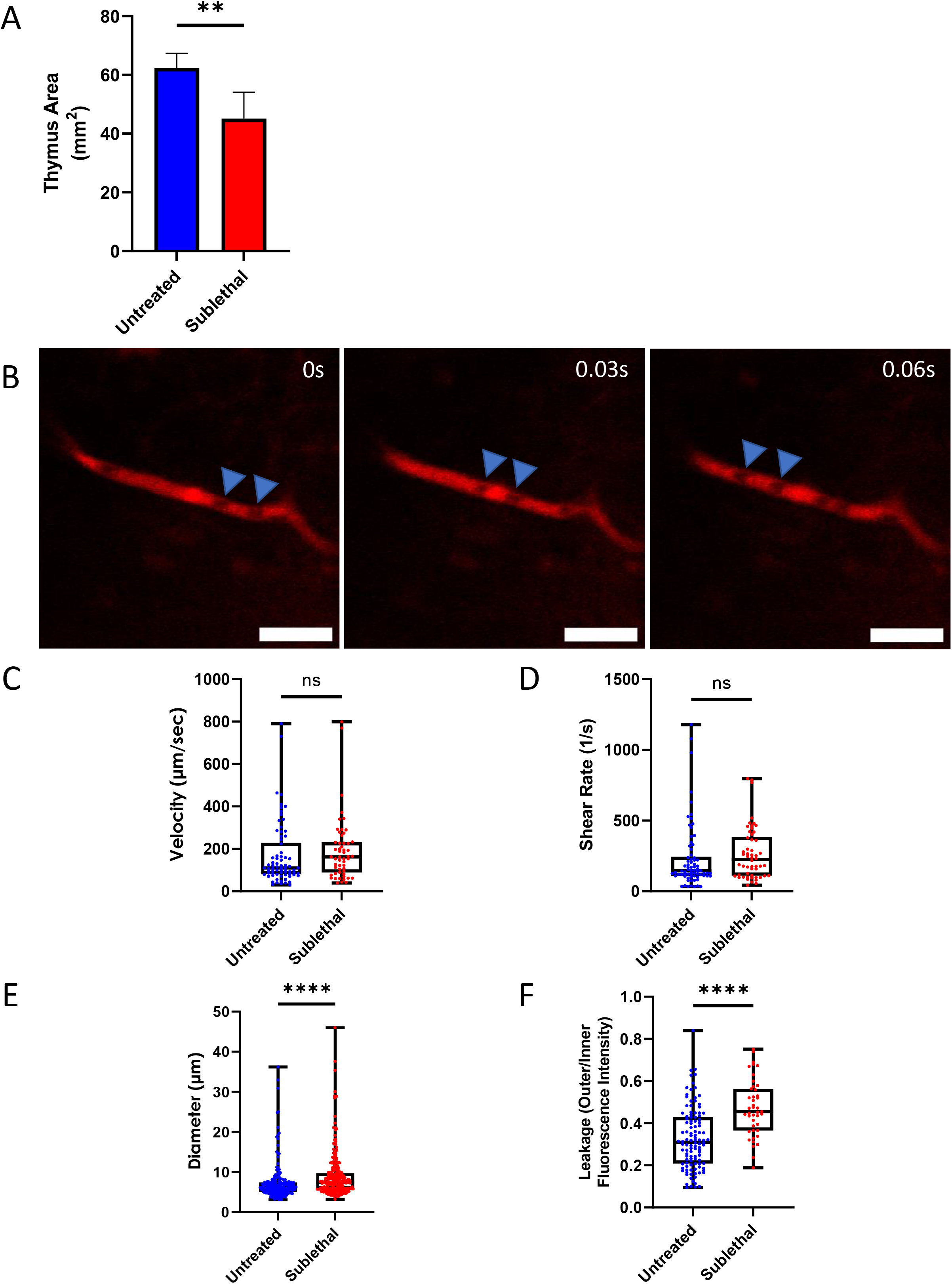
In Vivo Comparison of Native Thymus Vasculature in SL-TBI and Untreated Mice. Quantification of thymus area **(A)** from untreated and SL-TBI mice. **(B)** Example negative contrast labeled blood flow in the SL-TBI thymus from a UBC-GFP mouse showing the movement of RBCs through blood vessels over time. Red: blood vessels (Evans Blue); Blue Arrows: negative contrast labeled RBCs. (Scale bars ~ 25 μm). Quantification of blood flow velocity **(C)**, vessel shear rate **(D)**, vessel diameter **(E)**, and vessel leakage **(F)** in the thymus of untreated and SL-TBI UBC-GFP mice.

### Ex Vivo Imaging of Cleared Thymus Tissue

To validate our in vivo data showing anatomical changes to the native thymus blood vessel network, we imaged optically cleared whole thymus lobes (n = 3 mice each for SL-TBI treated and untreated controls) using a modified ultimate 3D imaging of solvent-cleared organs (uDISCO) protocol, where the sample is chilled for the entirety of the imaging session (Pan et al., 2016; Qi et al., 2019). Visualization of the whole cleared thymus using two-photon microscopy 24 hrs after SL-TBI revealed anatomical changes to the blood vessel architecture when compared to the untreated mice (**Figs. 4A-D and Supp. Movies 6-7**). To quantify the changes to the thymic vascular network, we measured the blood vessel diameter in SL-TBI treated and untreated mice and found a significant increase in diameter in the SL-TBI mice within the cortex (mean blood vessel diameter = 7.0 and 6.3 μm, respectively; p < 0.0001; **Fig. 4E**). This statistically significant difference in blood vessel diameter was observed in both in vivo and ex vivo imaging of the thymus and no significant difference was observed between in vivo and ex vivo diameter measurements (p > 0.9999, data not shown). Despite this, we did not observe a significant difference (p = 0.3497) in thymus cortical vessel frequency (defined as the number of vessel segments per mm^2^) between SL-TBI treated and untreated mice (vessel frequency = 514.2 and 482.9 segments/mm^2^, respectively; **Fig. 4F**). Further analysis also revealed a significant increase (p < 0.0001) in thymus cortical blood vessel area (defined as the number of vessel pixels in a FOV over the total number of pixels in the image) between the SL-TBI treated and untreated mice (mean % vessel area = 25.5% vs. 19.3%, respectively; **Fig. 4G**). Taken together, these results confirm that damage to the thymus vascular network occurs shortly after irradiation and that even if the overarching vascular network superstructure remains, the relative vascular volume increases due to shrinkage of the extravascular space and a small increase in vessel diameter.

**Figure 4.**
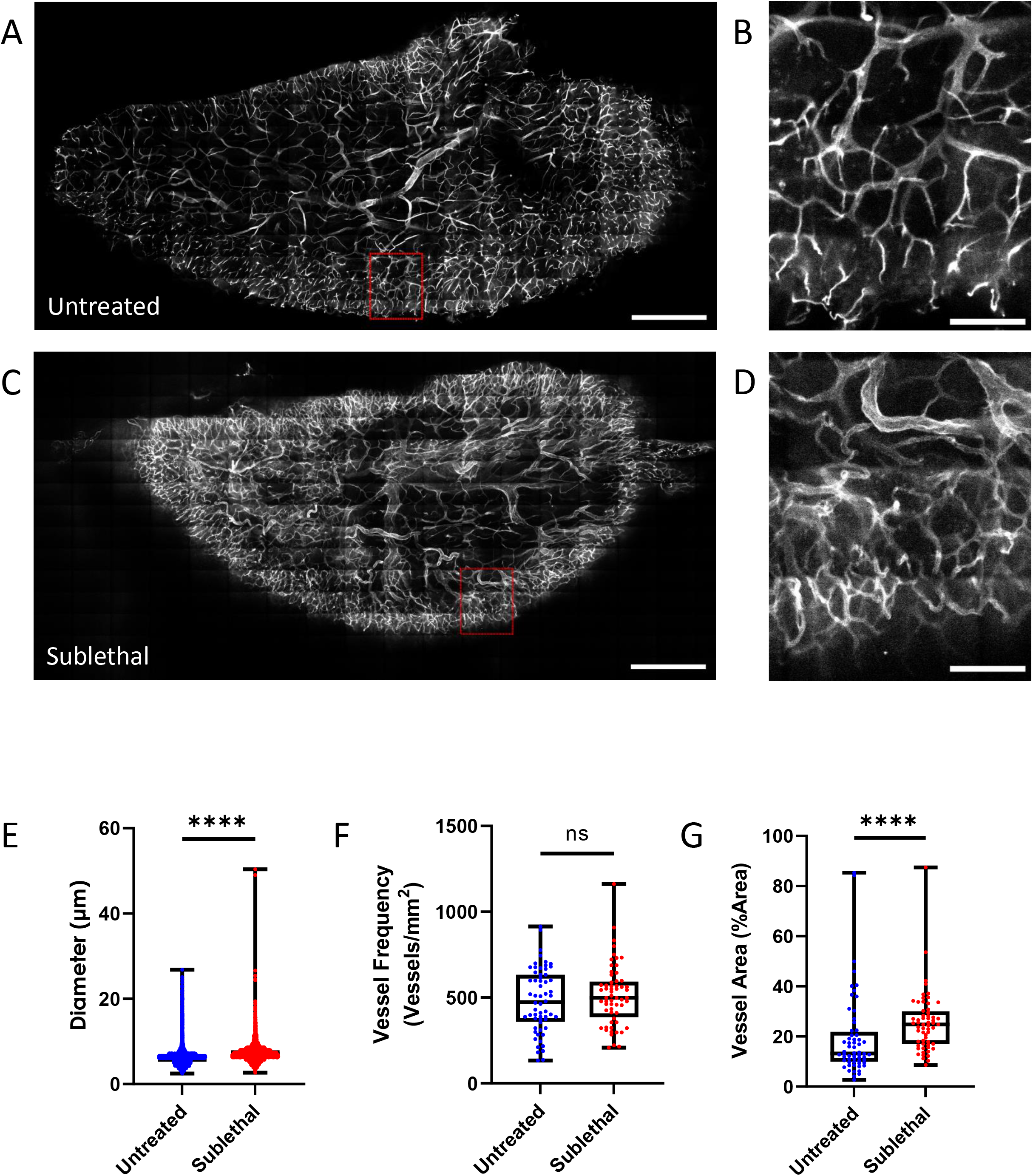
Changes to the Thymus Vasculature Observed Ex Vivo. **(A**) Representative average intensity projection of the untreated thymus. Grey: blood vessels (labeled with Alexa647 conjugated antibodies against CD31, CD144, and Sca-1); Red Box: cropped FOV. (Scale bar ~ 500 μm). **(B)** Cropped FOV of the untreated thymus. Grey: blood vessels (labeled with Alexa647 conjugated antibodies against CD31, CD144, and Sca-1). (Scale bar ~ 100 μm). **(C)** Representative average intensity projection of the SL-TBI thymus. Grey: blood vessels (labeled with Alexa647 conjugated antibodies against CD31, CD144, and Sca-1); Red Box: cropped FOV. (Scale bar ~ 500 μm). **(D)** Cropped FOV of the SL-TBI thymus. Grey: blood vessels (labeled with Alexa647 conjugated antibodies against CD31, CD144, and Sca-1). (Scale bar ~ 100 μm). Quantification of blood vessel diameter **(E)**, blood vessel frequency **(F)** and blood vessel area **(G)** in the untreated and SL-TBI thymus ex vivo.

## Discussion

Studies of T cell development and thymus regeneration have typically relied on techniques such as flow cytometry, immunohistochemistry, transplantation, and ex vivo imaging (Campinoti et al., 2020; Dzhagalov et al., 2012; Ladi et al., 2008; Le Borgne et al., 2009; Nakagawa et al., 2012; Witt et al., 2005). While these techniques provide valuable information about thymus biology and T cell development, only direct intravital visualization of the native thymus enables accurate characterization of the spatiotemporal dynamics of blood flow and maintains the natural chemical mileau within the thymus. Therefore, we developed a novel intravital imaging technique to surgically access the thymus and directly visualize it with two-photon microscopy. We were able to record stable videos and images within the native thymus and measure blood flow velocity, blood vessel diameter, shear rate and leakage in the thymus cortex. Due to imaging depth limitations incurred from light scattering and absorption, we imaged only the first ~150 μm of the native thymus, limiting observation mostly to the cortex. Nevertheless, we observed several larger blood vessels greater than 10 μm (up to 45 μm) in diameter deep within the thymus. As blood vessels in the thymus cortex tend to be smaller than in the medulla (Anderson et al., 2000; Kato, 1997; Raviola & Karnovsky, 1972), this observation potentially indicates the ability to image the thymus corticomedullary junction and even into the medulla with this technique. Further work will help to elucidate this finding.

While alterations to the thymus vascular structure have been reported 4 days after SL-TBI (Wertheimer et al., 2018), to our knowledge, no investigation has observed the effects of SL-TBI on native thymus hemodynamics. Using our imaging technique, we were able to study changes to hemodynamics in the thymus within 24 hours of SL-TBI. Although we observed a subtle but significant increase in blood vessel diameter and leakage shortly after SL-TBI, we found no significant difference in either blood flow velocity or shear rate. This data suggests that although the blood-thymus barrier has been compromised, as indicated by increased leakage, the blood flow and shear rate are largely unaffected at this timepoint. These results raise several questions regarding the role hemodynamics and blood vessel integrity may play in promoting thymus regeneration, such as creating a more open blood-thymus barrier for ETP homing and transmigration, and the abnormal exposure of thymocytes and stromal cells to blood plasma constituents. Intravital study of the blood-thymus barrier at later time points would likely provide additional information about the role it plays in supporting ETP homing and cellular expansion, but stress-induced thymus shrinkage (Wertheimer et al., 2018) increases the difficulty of imaging at later timepoints after SL-TBI. Future technical innovations such as endoscopic imaging (Jung et al., 2013; Kim et al., 2012; Vinegoni et al., 2012) and improved stabilization techniques (Lee et al., 2012; Vinegoni et al., 2015) may allow for the study of the thymus vasculature system and microenvironment across the entire thymus recovery period.

In addition, we utilized a modified tissue clearing method (Pan et al., 2016) combined with whole organ ex vivo two-photon imaging to validate the observed changes to the blood vessel network following SL-TBI. We characterized the thymic blood vascular system and quantified changes to the vessel diameter, volume, and frequency. Our imaging results revealed dramatic changes to the overall structure of the blood vessel network with a significant increase in vessel diameter and area. Despite this, there were no significant differences in vessel frequency in SL-TBI thymi compared to the untreated control group. Coupled with the observation that the thymus undergoes significant shrinkage after SL-TBI, these results are in line with previous studies that showed the thymus vasculature is relatively radioresistant (Wertheimer et al., 2018).

Overall, this study demonstrates a powerful new imaging method to directly investigate the native thymus and enables the characterization of the thymic microenvironment in ways not previously deemed possible. We demonstrated that shortly after SL-TBI, the thymus vascular structure undergoes rapid and significant changes which may be of clinical relevance in the context of thymus regeneration and immune system recovery following HCT.

## Acknowledgments

We would like to thank Drs. Jennifer Manilay and Katrina Hoyer of UC Merced for their very helpful input, suggestions, and encouragement throughout this work. We would like to thank Mr. Mario Muniz for his help in designing and testing the sample cooler system. We would like to thank Mr. Reagan Chan for his assistance with cleared thymus sample analysis. We would like to thank the staff of the Department of Animal Research Services at UC Merced for their continued assistance with the breeding and maintenance of our animal colony. We would like to thank Drs. Karin Gustafsson and David Scadden of Harvard University and Massachusetts General Hospital for encouraging us to develop intravital imaging of the native thymus. Schematic figures were created with BioRender.com. This work was funded by faculty startup funds provided to JAS by the University of California and through support of the NSF-CREST: Center for Cellular and Biomolecular Machines at the University of California, Merced (NSF-HRD-1547848 and NSF-HRD-2112675).

## Author Contributions

NS and CSB designed and performed experiments, analyzed and interpreted data and wrote the manuscript. RV and VO performed image analysis under supervision of CSB. NA helped develop the tissue clearing protocol. JAS conceived and supervised the project, interpreted data, and wrote the manuscript.

