## Supplementary Information for "Intravital Two-Photon Microscopy of the Native Thymus"

### Resource Availability

#### Lead Contact

#### Materials Availability

This study did not generate new unique reagents.

#### Data and Code Availability

All data was analyzed with standard programs and packages, as detailed in the Experimental Methods section. Software scripts and 3D modeling links are detailed in the Experimental Methods section. Image data is available upon request.

### Experimental Methods

#### Experimental Animals

Male and Female UBC-GFP (C57BL/6-Tg (UBC-GFP)30Scha/J) mice between the age of 8-12 weeks were used for intravital imaging experiments and wild-type (C57BL/6J) mice were used for ex vivo imaging experiments. Mice were housed under pathogen-free conditions in the University of California, Merced's vivarium with autoclaved feed and water, and sterile microisolator cages. The University of California Merced Institutional Animal Care and Use Committee (IACUC) approved all animal work. All mice were euthanized in accordance with IACUC-approved methods after imaging and before tissue collection.

#### Animal Irradiation

Mice received SL-TBI from a Precision X-Rad 320 at a dose of 4.5 Gy ~ 24hrs before intravital imaging or tissue extraction.

#### Preparation of Thymus for Ex Vivo Whole Organ Imaging

Mice were anesthetized with isoflurane (induction 3%, maintenance 1.5%, 100% $O_2$ at 1L/min). Fluorescent antibodies (anti-mouse Ly-6A (Sca-1) A647 Biolegend 108118; anti-mouse CD144 (VE-cadherin) A647 Biolegend 138006; and anti-mouse CD31 A647 Biolegend 102516) were retro-orbitally injected to label blood vessels. 20 minutes after injection, animals were perfused intracardially with phosphate buffered saline (1X), followed by cold paraformaldehyde (4%, pH 7.4) for 5–10 min. The dissected thymus was postfixed in paraformaldehyde (4%) overnight at 4°C. To optically clear the thymus, we modified the uDISCO tissue clearing protocol by keeping the sample at 4°C for the entirety of the imaging session (Pan et al., 2016). Postfixed thymus lobes were dehydrated with tert-butanol (Sigma-Aldrich; SHBM5332). Dehydration solutions were prepared by mixing tert-butanol and distilled water at various concentrations (30%/50%/70%/90%/100% tert-butanol). Next, the thymus was incubated in dichloromethane (DCM; Sigma-Aldrich, SHBJ8352) for the delipidation process. The tissue was then placed in BABB-D4, prepared by mixing BABB (benzyl alcohol + benzyl benzoate 1:2, Sigma-Aldrich; 24122 and W213802) with diphenyl ether (DPE; Alfa Aesar, A15791) at a ratio of 4:1 and adding 0.4% vol DL-alpha-tocopherol (Vitamin E; Alfa Aesar, A17039). Cleared thymus lobes were mounted in a custom-designed slide well filled with BABB-D4 and sealed with solvent-resistant silicone sealant (DOWSIL™ 730).

#### Animal Preparation for Intravital Thymus Imaging

Mice were initially anesthetized (3-4% Isoflurane for induction, 1.5-2% for maintenance, 100% $O_2$ at 1L/m) and then mounted to a heating pad to maintain normal body temperature. The right thoracic and abdominal regions were shaved, and the skin cleaned with 70% alcohol wipes. The mouse was then intubated using a laryngoscope and a 22-G angiocatheter (Exel Int, 26746). An injection of ketamine (100mg/kg) / xylazine (15mg/kg) was administered via IP injection before the thoracotomy. To perform the thoracotomy, an incision was made through the 2$^{nd}$ intercostal space above the thymus and between the internal thoracic vein and sternum and expanded by inserting rib retractors into the intercostal space until the thymus was visible. A cauterizer was used to stop any excessive bleeding around the surgical site. The mice were euthanized after intravital imaging and the thymi were removed for ex vivo thymus imaging.

#### Thymus Holder Placement

The adhesion stabilization holder consists of a stabilization ring with an inner diameter of 2.5 mm attached to an angled stabilization arm and was modeled in Openscad and 3D printed in polylactic acid (PLA). The holder was directly attached to the exposed thymus via a tissue safe adhesive (Vetbond, 084-1469SB) applied to the underside of the stabilization ring via a small paintbrush. After attachment, warm methyl cellulose (Sigma-Aldrich, M0512) was applied to the center of the holder to act as an immersion fluid for imaging.

#### Two-Photon Microscopy

Imaging was performed with a custom-built two-photon video-rate microscope (Bliq Photonics) equipped with two femtosecond lasers (Insight X3 and MaiTai eHP DS, Spectra Physics). During intravital imaging, the Insight X3 and Maitai laser wavelengths were tuned to 800 nm and 950 nm, respectively, and for ex vivo imaging the Insight X3 and Maitai laser wavelengths were tuned to 820 nm and 1220 nm, respectively. Three fluorescent channels were acquired (503-538 nm, 572-608 nm, and 659-700 nm). For two-photon imaging, a 25x water immersion objective (Olympus; XLPLN25XWMP2) with a 1.05 numerical aperture was used to record video at 30/60/120 frames per second at a resolution of ~0.31 µm/pixel. To label thymus blood vessels in vivo, Evans's blue was injected retro-orbitally after the thymus holder was placed.

#### Image Analysis

For image processing and measurements of blood vessel diameter, shear rate, leakage, frequency and area analysis, and manual blood flow quantification, Fiji (ImageJ 1.53f51) was used. All ImageJ scripts used for analysis and thymus adhesion holder 3D models are available online (https://github.com/SpencerLab-BIO/NativeThymusScripts). Image brightness/contrast was adjusted in Fiji for display purposes and images were cropped to remove vignetting. A previously published MATLAB (2020a) script was used to calculate blood flow automatically (Wu et al., 2021). Cleared thymus datasets were stitched together using the Grid Stitching plugin in Fiji.

During intravital imaging, videos of blood flow in the thymus were recorded. Blood flow velocity was calculated with two different methods depending on tissue stability during imaging. When minimal tissue movement was present, we used pLSPIV to calculate blood flow velocity automatically (Kim et al., 2012; Wu et al., 2021). However, when significant tissue movement was present, blood flow velocity was calculated manually by tracking the change in the position of individual RBCs over time. Video frames were first aligned via the Linear Stack Alignment with SIFT plugin in Fiji and then the displacement of the approximate RBC centroid between frames was used to calculate blood flow per blood vessel.

Blood vessel diameters both in vivo and ex vivo were calculated via two different methods depending on the signal-to-noise ratio and the orientation of the blood vessel relative to the focal plane. When high contrast was observed between blood vessels and background, and vessels were orientated parallel to the focal plane, a modified version of the VasoMetrics2 (McDowell et al., 2021) Fiji script was used. The only modifications made to the VasoMetrics2 script were done to improve user experience and the function related to vessel diameter calculation was not modified. Alternatively, when there was poor contrast between the vessels and background or vessels were orientated perpendicular to the focal plane, vessel diameter was measured manually via the Straight-line tool in Fiji. Blood vessel leakage measurements were taken at least 10 min after Evans blue injection by averaging 30 frames of blood vessel footage. Leakage was calculated as the ratio between the fluorescent intensity inside of a blood vessel vs. outside immediately adjacent to the blood vessel. The shear rate for individual vessels was approximated as (8 × velocity) / diameter as previously described (Bixel et al., 2017).

To measure blood vessel diameter, frequency, and area in the optically cleared thymus, 3D datasets of the thymus vasculature were downscaled by half, the Despeckle and Gaussian Blur ($\sigma = 2$) filters were applied, and the brightness/contrast was adjusted in Fiji. A mask of the outline of the thymus was then generated by downscaling the thymus by half and then either using a custom weka model generated by the Labkit plugin (Arzt et al., 2022) or manually tracing the perimeter of the thymus. The mask of the thymus was then upscaled by 2 and a Euclidian Distance Map of the thymus was generated with the 3DSuite plugin (Ollion et al., 2013). The resulting distance map was thresholded to generate a mask corresponding to a volume ~150 µm from the edge of the thymus. To quantify blood vessel diameter and density in the cleared thymus, twenty random non-overlapping 300 x 300 µm FOV whose centroid was <150 µm from the edge of the thymus were used. The previously mentioned ~150 µm mask was used to limit the center of the FOV to within 150 µm from the edge of the thymus. Vessel diameter was calculated as previously described and vessel frequency was calculated as the number of vessels in the FOV divided by the area of the thymus in the FOV. The vessel area was calculated by thresholding the blood vessels in each FOV via the Otsu method and then calculating the % area in each FOV occupied by blood vessels.

To measure the thymus volume, the dissected thymus was placed on a gridded reference and the % area of the grid covered by the thymus was calculated (Wan et al., 2018).

#### Statistical Analysis

The Mann-Whitney U-test and Student's t-tests were performed in GraphPad Prism to test for statistical significance, depending on whether datasets were normally distributed. A p-value $< 0.05$ was considered statistically significant (Data are presented as mean ± SD; *p<0.05, **p<0.01, ***p<0.001; ****p<0.0001).

### Experimental Methods References

Arzt, M., Deschamps, J., Schmied, C., Pietzsch, T., Schmidt, D., Tomancak, P., Haase, R., & Jug, F. (2022). LABKIT: Labeling and Segmentation Toolkit for Big Image Data. *Frontiers in Computer Science*, *4*. https://www.frontiersin.org/articles/10.3389/fcomp.2022.777728

Bixel, M. G., Kusumbe, A. P., Ramasamy, S. K., Sivaraj, K. K., Butz, S., Vestweber, D., & Adams, Ralf. H. (2017). Flow Dynamics and HSPC Homing in Bone Marrow Microvessels. *Cell Reports*, *18*(7), 1804–1816. https://doi.org/10.1016/j.celrep.2017.01.042

Kim, T. N., Goodwill, P. W., Chen, Y., Conolly, S. M., Schaffer, C. B., Liepmann, D., & Wang, R. A. (2012). Line-Scanning Particle Image Velocimetry: An Optical Approach for Quantifying a Wide Range of Blood Flow Speeds in Live Animals. *PLOS ONE*, *7*(6), e38590. https://doi.org/10.1371/journal.pone.0038590

McDowell, K. P., Berthiaume, A.-A., Tieu, T., Hartmann, D. A., & Shih, A. Y. (2021). VasoMetrics: Unbiased spatiotemporal analysis of microvascular diameter in multi-photon imaging applications. *Quantitative Imaging in Medicine and Surgery*, *11*(3), 969–982. https://doi.org/10.21037/qims-20-920

Ollion, J., Cochennec, J., Loll, F., Escudé, C., & Boudier, T. (2013). TANGO: A generic tool for high-throughput 3D image analysis for studying nuclear organization. *Bioinformatics*, *29*(14), 1840–1841. https://doi.org/10.1093/bioinformatics/btt276

Pan, C., Cai, R., Quacquarelli, F. P., Ghasemigharagoz, A., Lourbopoulos, A., Matryba, P., Plesnila, N., Dichgans, M., Hellal, F., & Ertürk, A. (2016). Shrinkage-mediated imaging of entire organs and organisms using uDISCO. *Nature Methods*, *13*(10), Article 10. https://doi.org/10.1038/nmeth.3964

Wan, P., Zhu, J., Xu, J., Li, Y., Yu, T., & Zhu, D. (2018). Evaluation of seven optical clearing methods in mouse brain. *Neurophotonics*, *5*(3), 035007. https://doi.org/10.1117/1.NPh.5.3.035007

Wu, J. W., Jung, Y., Yeh, S.-C. A., Seo, Y., Runnels, J. M., Burns, C. S., Mizoguchi, T., Ito, K., Spencer, J. A., & Lin, C. P. (2021). Intravital fluorescence microscopy with negative contrast. *PLOS ONE*, *16*(8), e0255204. https://doi.org/10.1371/journal.pone.0255204

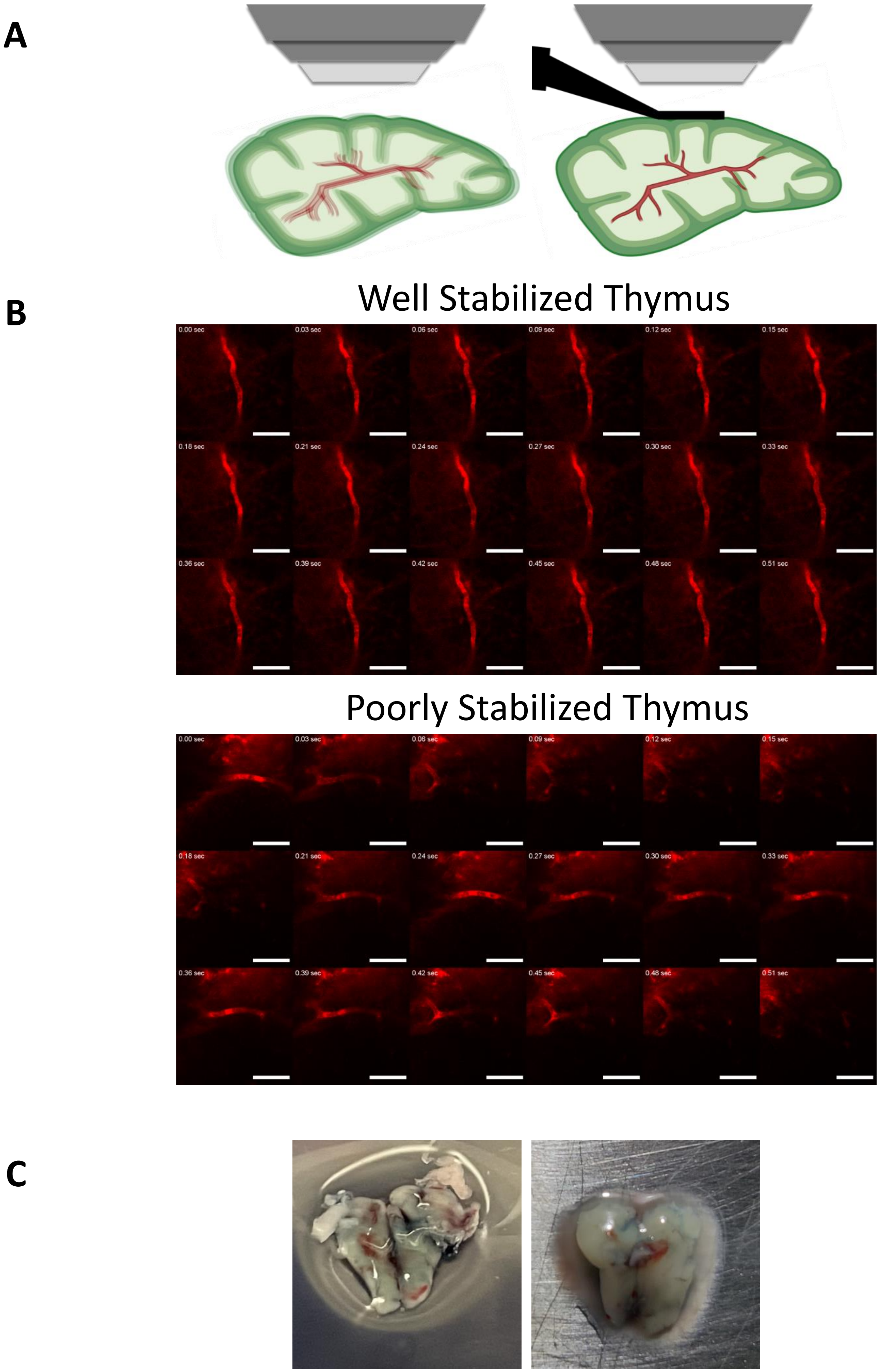


Supplementary Figure 1

A

B

C

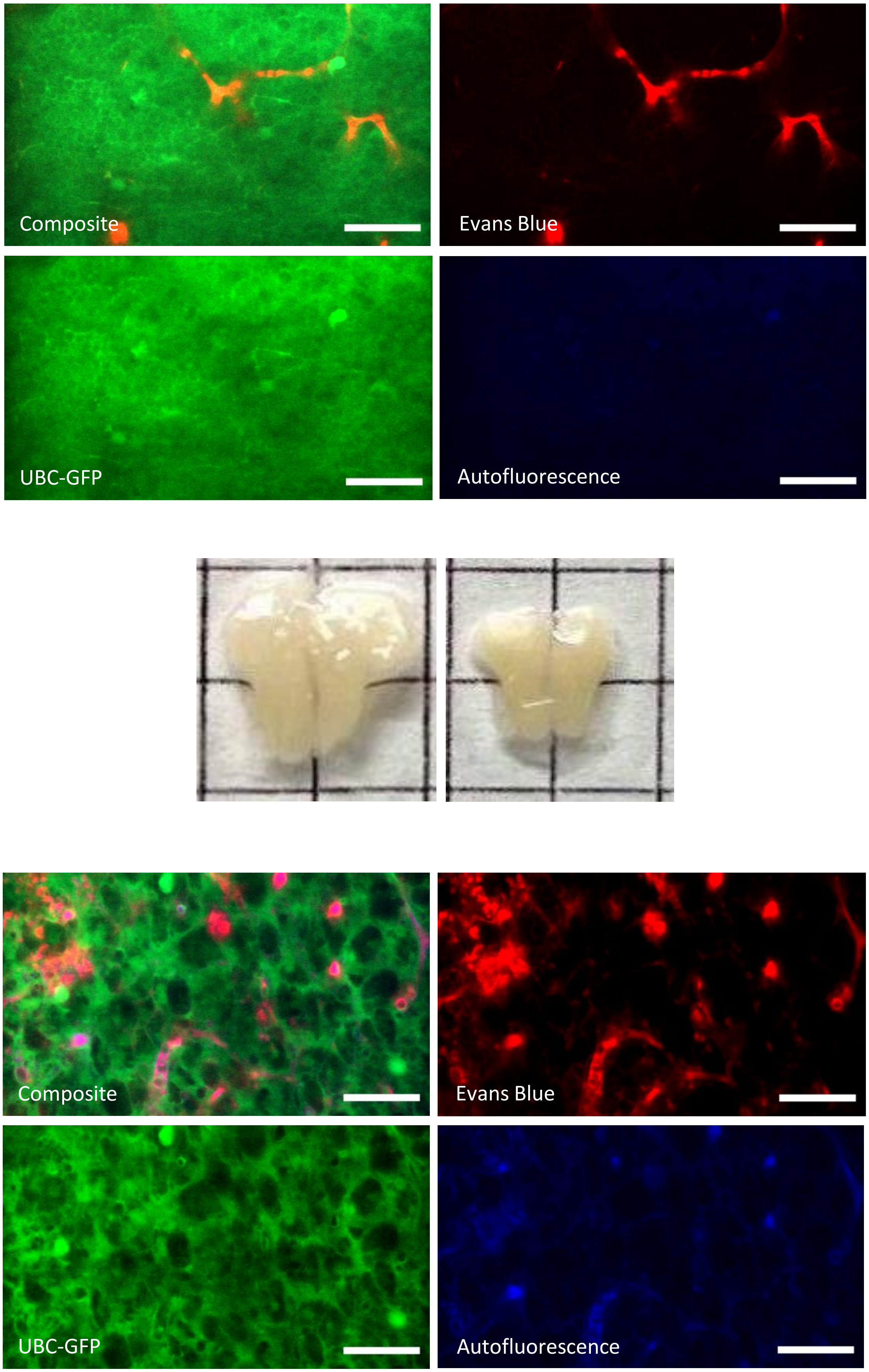


Supplementary Figure 2

### Supplementary Figure and Movie Legends

**Supplementary Figure 1. Thymus Stabilization via an Adhesion Holder**

**(A**) Representative diagram of an unstabilized (left) and stabilized (right) thymus during imaging. Red: blood vessels/thymus capsule; Green: GFP; Black: adhesion holder.

**(B)** Representative montages of blood vessels in well- and poorly-stabilized thymi. Red: blood vessels (Evans blue). (Scale bar ~ 50 µm).

**(C)** Representative images of dissected thymi from untreated (left) and SL-TBI (right) mice after receiving an Evans blue injection, demonstrating successful perfusion of the thymus.

**Supplementary Figure 2. Dissected and Ex Vivo Thymus**

**(A)** Representative average intensity projection of the thymus from an untreated mouse ex vivo. Red: blood vessels (Evans blue); Green: GFP; Blue: autofluorescence. (Scale = 50um).

**(B)** Representative size difference of thymi from untreated (left) and SL-TBI (right) mice. (Black square border ~ 5 mm)

**(C)** Representative average intensity projection of the thymus from a SL-TBI mouse ex vivo. Red: blood vessels (Evans blue); Green: GFP; Blue: autofluorescence. (Scale bar ~ 50 µm).

**Supplementary Movie 1: Representative Intravital Two-Photon Zstack of the Native Thymus.** Representative intravital two-photon zstack of the native thymus in a 10 week old UBC-GFP mouse. Red: blood vessels (Evans blue); Green = GFP; Blue = autofluorescence. (Scale bar ~ 50 µm, Zstep size = 2 µm)

**Supplementary Movies 2/3: Representative Blood Flow within the Untreated Native Thymus**

**Supplementary Movie 2:** Representative video showing blood flow of the thymus in vivo in an untreated 10 week old UBC-GFP mouse. Red: blood vessels (Evans blue); Green = GFP; Blue = autofluorescence. (Scale bar ~ 50 µm)

**Supplementary Movie 3:** Red channel only video corresponding to Movie 2. Grey: blood vessels (Evans blue). (Scale bar ~ 50 µm)

**Supplementary Movies 4/5: Representative Blood Flow within the SL-TBI Native Thymus**

**Supplementary Movie 4:** Representative video showing blood flow of the thymus in vivo in a SL-TBI 10 week old UBC-GFP mouse. Red: blood vessels (Evans blue); Green = GFP; Blue = autofluorescence. (Scale bar ~ 50 µm)

**Supplementary Movie 5:** Red channel only video corresponding to Movie 4. Grey: blood vessels (Evans blue). (Scale bar ~ 50 µm)

**Supplementary Movies 6/7: 3D Model of the Optically Cleared Thymus Vasculature**

**Supplementary Movie 6:** Representative 3D model of the optically cleared thymus vasculature from an untreated mouse. Grey = blood vessels (labeled with Alexa647 conjugated antibodies against CD31, CD144, and Sca-1). (Scale bar ~ 250 µm)

**Supplementary Movie 7:** Representative 3D model of the optically cleared thymus vasculature from a SL-TBI mouse. Grey = blood vessels (labeled with Alexa647 conjugated antibodies against CD31, CD144, and Sca-1). (Scale bar ~ 250 μm)
